# Visual analysis of spatial transcriptomics data with RedeViz

**DOI:** 10.1101/2024.07.09.602652

**Authors:** Dehe Wang, Xianwen Ren

## Abstract

Spatial transcriptomics (ST) technologies are powerful tools to illustrate the spatial hierarchy and heterogeneity of tissues with the lens of multiplexed gene readouts. However, ST technologies generate sequence data rather than images, preventing intuitive examination of the cellular contexture of tissues. Moreover, the inherent sparsity of ST data caused by molecular crowdedness and sequencing dropouts poses great challenges to accurate and clear visualization. In this study, we introduce RedeViz, a toolkit crafted for enhancing and visualizing subcellular-resolution ST data. RedeViz applies a pixel-level enhancement strategy, visualizes ST data in automatic or customized manners, and can display the cellular and genic spatial patterns with effects akin to HE staining. Strict evaluations confirm that RedeViz fits a wide range of ST platforms, including Xenium, Visium HD, MERFISH, CosMx, Stereoseq, as well as spatial proteomic platforms like CODEX. The impressive performance of RedeViz across various scales from cell-, tissue-, organ-, to organism-levels brings us a universal “What You See Is What You Get” framework for visual analysis of ST data.

## Introduction

Subcellular-resolution spatial transcriptomics (ST) technologies are powerful tools for understanding tissue archetecture^1– 3^ and disease abnormalities^4–6^ by capturing the spatial arrangement of gene expression patterns. Such technologies, including imaging-based methods (such as MERFISH^7^, CosMx^8^, and Xenium^9^) or next-generation sequencing (NGS)based methods (such as Visium HD^10^ and Stereo-seq^11^), hold the potential to intuitively visualize the spatial hierarchy and heterogeneity in biological structures.

However, the absence of clear cell boundaries and the inherent sparsity^14–16^ of ST data present challenges in accurately visualizing the data, limiting the characterization of fine spatial patterns of cells and genes. Current visualization methods often rely on manual annotation of cell types and assignment of colors for visualization^4–6,12,13,17^, leading to overlooks of important cellular structures, potential inaccuracies, or even artifacts due to issues introduced by manual manipulation. On one hand, coarse-grained visualization methods, including staining grids^5,6,18–20^ or predicted cells^4,5,17,21^, often lead to loss of intricate details due to resolution reduction. On the other hand, raw signal is now not applicable to visualize the spatial distribution of gene expression due to the signal sparseness of current ST data^12,22,23^. An automatic resolution-preserving method with signal enhancement is thus urgently needed.

Inspired by previous works that researchers mapped all spectral characteristics of mass spectrometry to color space and automatically visualized mass spectrometry imaging (MSI) data at spot resolution^24^, we introduce RedeViz, an unsupervised segmented-free method that can automatically visualize the cellular and genic spatial patterns of ST data at pixel resolution. Analyses of Xenium, MERFISH, and CosMx datasets with tissue-level complexity demonstrate that RedeViz achieves visual effects comparable to hematoxylin-eosin (HE) and immunofluorescence (IF) images for automatic cell state and gene expression visualization tasks. RedeViz also shows strong performance on spatial proteomics (SP) platforms, such as CODEX. Using Visium HD datasets, RedeViz shows the ability to automatically visualize the cell contexture akin to HE images. The Stereo-seq embryo dataset showcases RedeViz’s performance for organism-level tasks of visualizing ST data. Furthermore, the targeted enhancement strategy enables RedeViz to perform computational anatomy of embryos, with more detailed cellular contexture of each organ illustrated automatically as exemplified by the fine structures of the kidney and inner ear. In summary, RedeViz shows the potential to serve as a universal framework for visual analysis of ST data with a What You See Is What You Get (WYSIWYG) effect.

## Results

### RedeViz: A workflow for enhancement and visualization of ST data

Here we develop Raising Dimension for Enhanced Visualization (RedeViz), a segmentation-free approach aimed at enhancing and visualizing subcellular-resolution ST data at pixel resolution (Supplementary Figure 1, and Methods). RedeViz involves two main steps: “raising dimension for enhancement” and “visualization of enhanced ST data”. In the “raising dimension for enhancement” step, RedeViz constructs a reference data to extract information like cell state, cell type, and gene expression. The reference data can be constructed from a single-cell RNA sequencing (scRNA-seq) dataset, an external ST dataset, or the ST dataset itself to be visualized. Recognizing that not all samples may have accessible matched scRNA-seq data, RedeViz offers the flexibility to utilize pre-segmented ST data as a reference in particular. It then enhances the sparse ST data by adding extra derived dimensions (dDims) from reference data by aligning the two datasets pixel by pixel. This process fills in gaps caused by data sparsity and imputes missing information to the ST data. Moving to the “visualization of enhanced ST data” step, RedeViz offers both fine-grained (pixel resolution) and coarsegrained visualization. In fine-grained visualization, RedeViz can visualize ST data using the enhanced dDims in an automatic manner or alternatively visualize the cell state and cell type distributions with user-customized colors. Moreover, RedeViz can visualize gene expression for genes not directly measured in the ST data because of the enhancement step, which can impute the ST data with the help of whole-transcriptome scRNA-seq data. In coarse-grained visualization, RedeViz can identify and visualize mesoscopic or macroscopic spatial domains. With these capabilities, RedeViz facilitates a comprehensive visualization of information encompassing various dimensions and levels of granularity within the ST data. This advancement brings it closer to the WYSIWYG effect for an ideal ST data visualization.

### Enhancing and visualizing ST data to achieve effects akin to HE staining

We used a breast cancer Xenium dataset^9^ to evaluate RedeViz’s performance at tissue-level complexity. Automatic cell state visualization using RedeViz revealed that it effectively captured the fine tissue structure within the ST data, compared to hematoxylin-eosin (HE) staining and manually annotated cell type distribution (Figure 1a-c). RedeViz accurately depicted structures such as immune-encapsulated ductal carcinoma in situ (DCIS), non-immune-encapsulated DCIS, and adipose tissues. Particularly, RedeViz captured the intricate structure of adipocytes with vascular endothelial cells embedded, matching results obtained through HE staining (Figure 1a, c). This suggests that RedeViz can visualize fine structures in image-based ST data with accuracy comparable to HE staining. In terms of cell type annotation, RedeViz proved accuracy across various structures, similar to its performance in automatic cell state visualization (Figure 1d). Compared with the raw Xenium signal and binning-based visualization, the imputed gene expression patterns of *CD3E* (T cells) and *VWF* (endothelial cells) closely matched the results of HE staining (Supplementary Figure 2a-d), indicating its ability to visualize global features while preserving high-resolution details. Moreover, RedeViz could impute genes not measured in the Xenium dataset (Supplementary Figure 2e), enabling discoveries with the help of wholetranscriptomics information.

**Figure 1.**
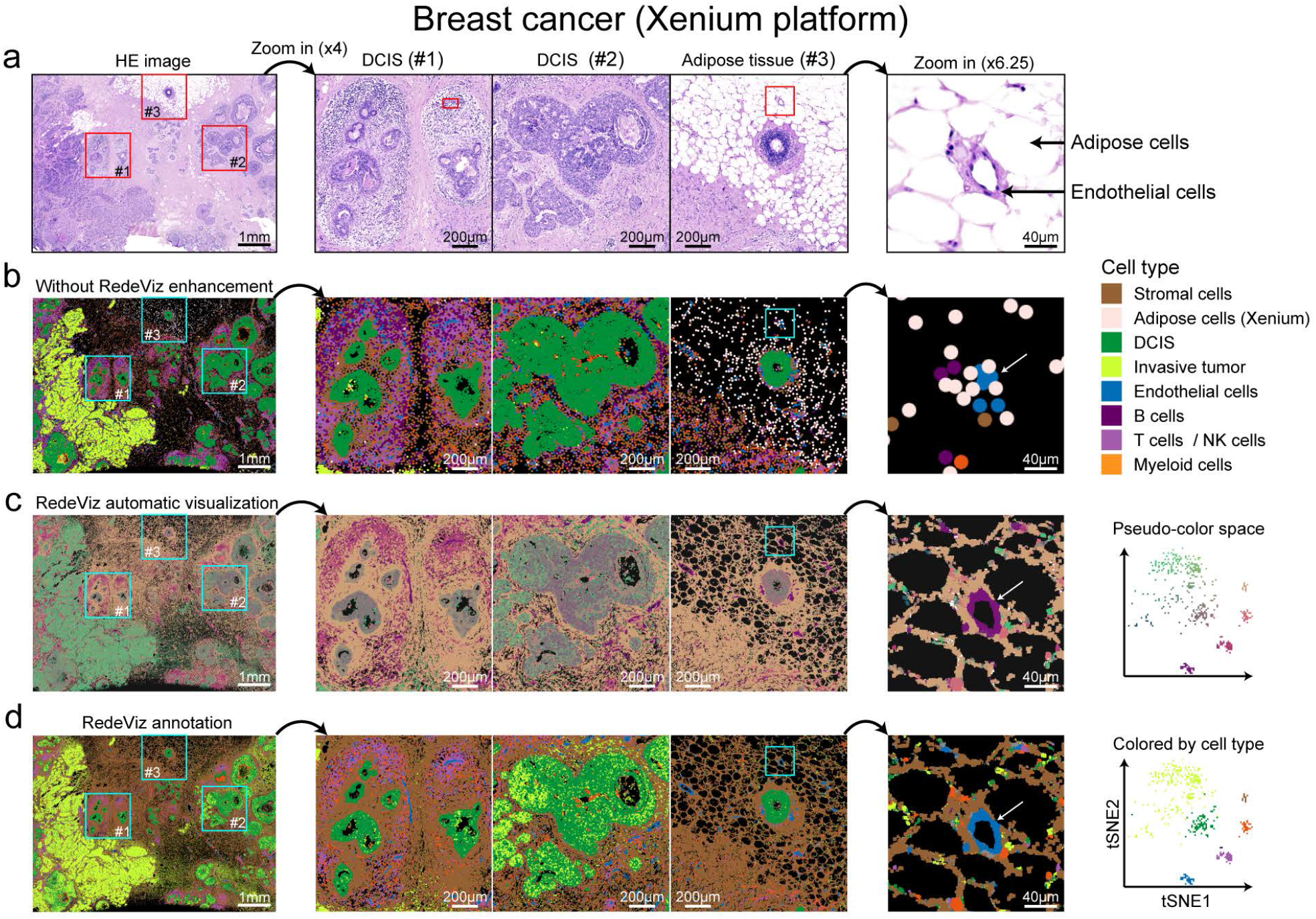
RedeViz visualization on breast cancer Xenium dataset. (a-d) The full region (left), three spatial structure (middle) and fine structure of adipose tissue (right) of HE image (a), manual cell type annotation (b), RedeViz automatic cell state visualization (c), and RedeViz annotation (d) after RedeViz enhancement are shown. The images of RedeViz automatic cell state visualization are colored by pseudo-color, and the images of manual cell type annotation and RedeViz cell type annotation are colored by cell types. And the vascular endothelial cells in fine structure of adipose tissue are indicated by arrows.

Subsequently, we applied RedeViz to enhance and visualize the mouse ileum MERFISH dataset^11^, and human nonsmall cell lung cancer (NSCLC) CosMx dataset^8^. In the mouse ileum, cells are organized in distinct layers. Starting from the basal side and progressing towards the villus side, the arrangement consists of myocyte cells, Paneth cells, stem cells, transit amplifying cells, and enterocyte cells (Figure 2a). RedeViz elucidates the layered arrangement of ileum cells after enhancement by the pre-segmented ST data (self-enhancement), no matter cell states or the cell types were visualized (Figure 2b-c). RedeViz also shows good performance by leveraging scRNA-seq as references (Supplementary Figure 3a-b). Additionally, RedeViz demonstrates its proficiency in imputing gene expression distributions for genes measured (*Mki67*) or unmeasured (*Alpi* and *Gm15293*) in the MERFISH dataset (Supplementary Figure 3c).

**Figure 2.**
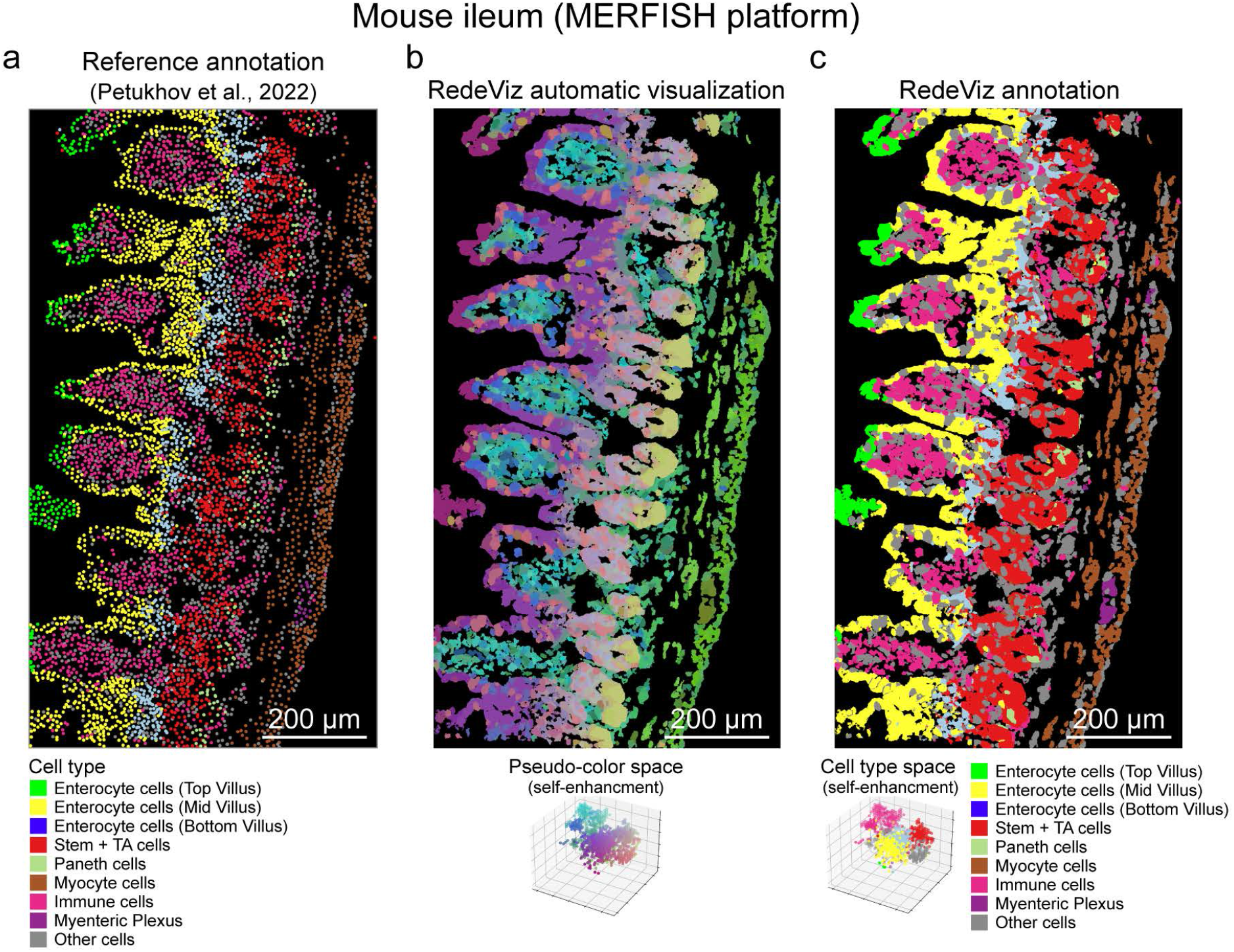
RedeViz visualization on mouse ileum MERFISH dataset. (a) The reference cell type annotation. (b-c) RedeViz automatic cell states visualization (b) and cell type annotation (c) after self-enhancement. The images of RedeViz automatic cell state visualization are colored by pseudo-color.

As for the NSCLC CosMx dataset (Figure 3a-b), RedeViz discriminates between tumor cells, normal epithelial cells, and immune cells in automatic cell state visualization (Figure 3c), ensuring precise cell type annotations (Supplementary Figure 4a). Notably, the annotated normal epithelial and tumor cells closely align with the epithelial cell marker (PanCK) positions in the immunofluorescence (IF) image (Figure 3d). RedeViz accurately predicts tumor cell locations within continuous tumor areas and identifies isolated cells away from tissue (Figure 3d), showcasing visual resolution comparable with IF. Moreover, compared to raw or binned signals, RedeViz displays the characteristics of genic spatial distribution while preserving high resolution (Supplementary Figure 4b-d).

**Figure 3.**
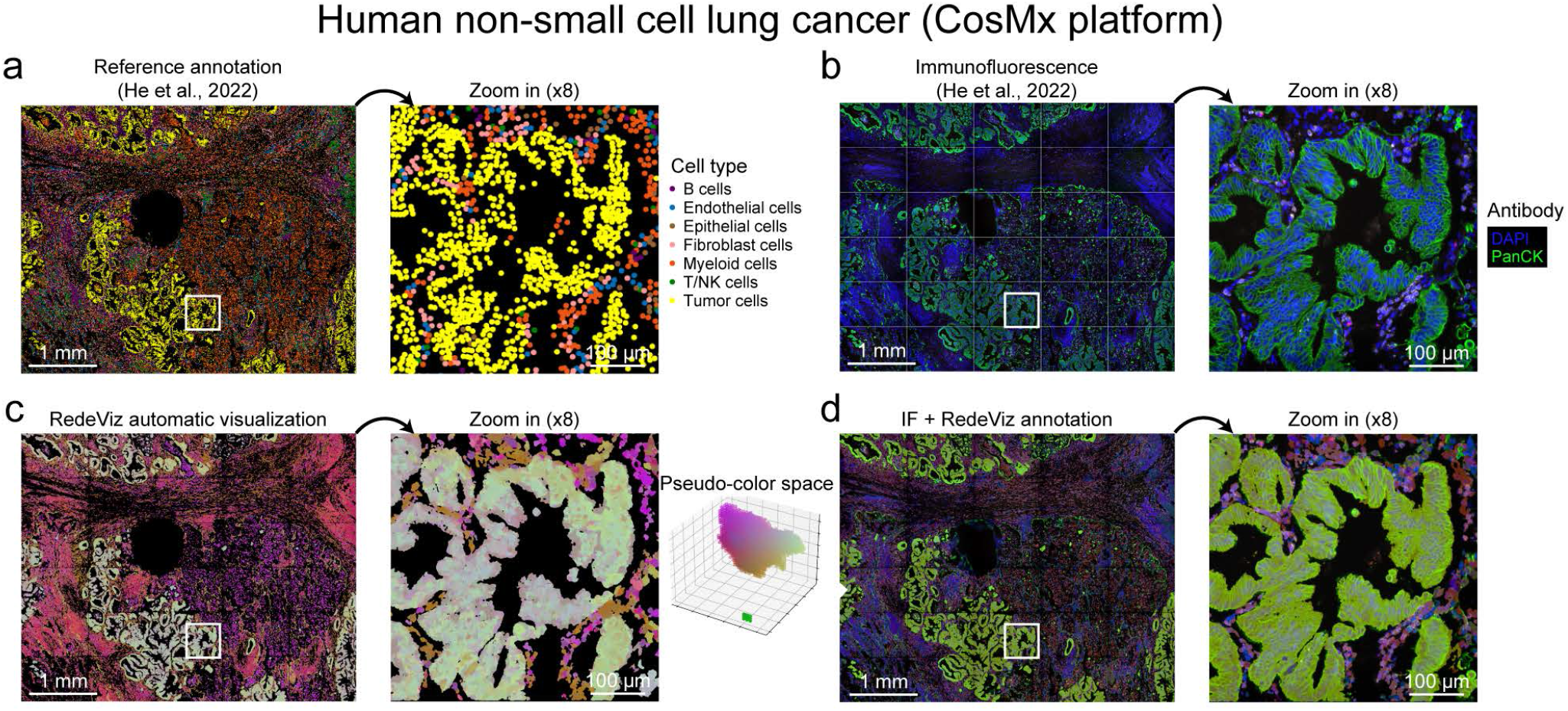
RedeViz visualization on human non-small cell lung cancer (NSCLC) CosMx dataset. (a) The reference cell type annotation. (b) The immunofluorescence (IF) of DAPI and PanCK. (c) RedeViz automatic cell state visualization. (d) Merged image of IF and RedeViz cell type annotation.

RedeViz can effectively visualize not only spatial transcriptomics data but also spatial proteomics data. We apply RedeViz to the human bone marrow CODEX dataset^13^. Compared to the reference annotation (Figure 4a), RedeViz effectively describes structures with weak DAPI signals (Figure 4b-c region #1), such as vascular smooth muscle cells (VSMCs). For locations with strong DAPI signals, RedeViz can also reveal the monolayer endothelial cell structure surrounded by T cells (Figure 4b-c region #2). Due to the lack of DAPI signal, the original visualization is not effective for structures such as VSMCs and plasma where cell nuclei are not easily captured (Figure 4a,c). After RedeViz enhancement, these structures can be visualized at pixel resolution (Figure 4b-d and Supplementary Figure 5). These outcomes highlight RedeViz’s adaptability across platforms and its ability to approximate a WYSIWYG visual effect at the tissue scale.

**Figure 4.**
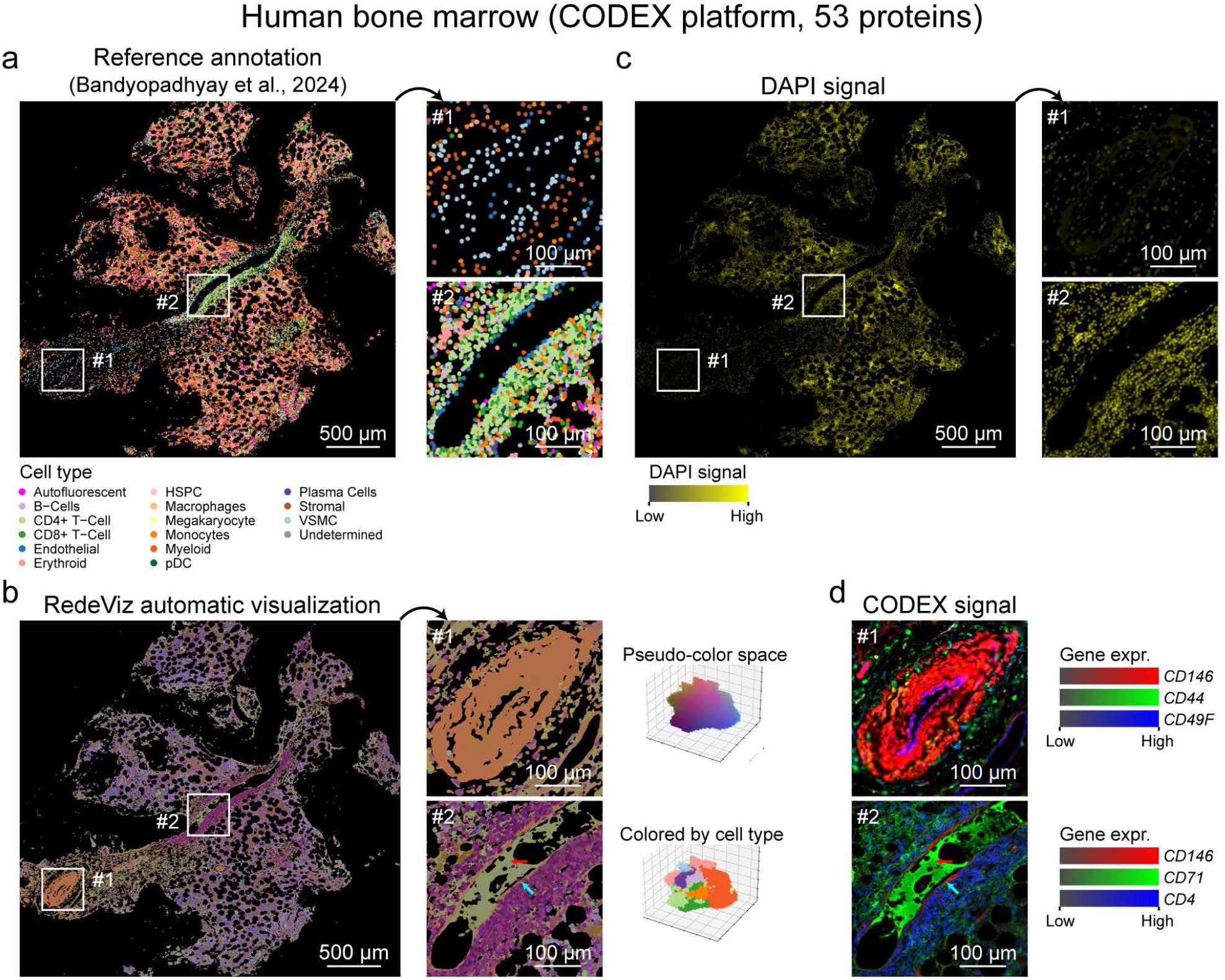
RedeViz visualization on human bone marrow CODEX dataset. (a) The reference cell type annotation. VSMC: vascular smooth muscle cells, HSPC: vascular smooth muscle cells. (b) RedeViz automatic cell state visualization. (c) DAPI signal. (d) CODEX signal of cell type specific proteins.

### Automagical visualization reveals hierarchical structures of mouse kidney

Next, we utilized the mouse kidney Visium HD dataset to assess RedeViz’s enhanced visualization capability at the organ level. Based on the HE staining, the kidney can be divided into four layers (Figure 5a). After self-enhancement, these four layers can be clearly visualized automatically based on the Visium HD data (Figure 5b). Automatic clustering based on the RedeViz visualization can result in 19 cell types, which were of distinct marker genes (Figure 5c). Therefore, RedeViz enhances the visual distinction of complex structures within the mouse kidney, which are difficult to distinguish with the original ST data or HE-stained images (Figure 5a-c). For the major cell types of L1-L4, RedeViz reveals both a clear hierarchical structure and a complex mosaic structure (Supplementary Figure 6a-b). The high consistency between the identified cell types and the corresponding marker positions confirms the accuracy of RedeViz. RedeViz also shows high accuracy for the minor cell types (Supplementary Figure 6c). Notably, we observed two closely attached layers of cells (*Acta2*^+^ and *Upk1b*^+^), where *Acta2* and *Upk1b* are marker genes for smooth muscle cells and urothelial cells^14^, respectively (Supplementary Figure 6d). The thickness of these two cell layers is only about 20μm, demonstrating the high resolution and accuracy of RedeViz.

**Figure 5.**
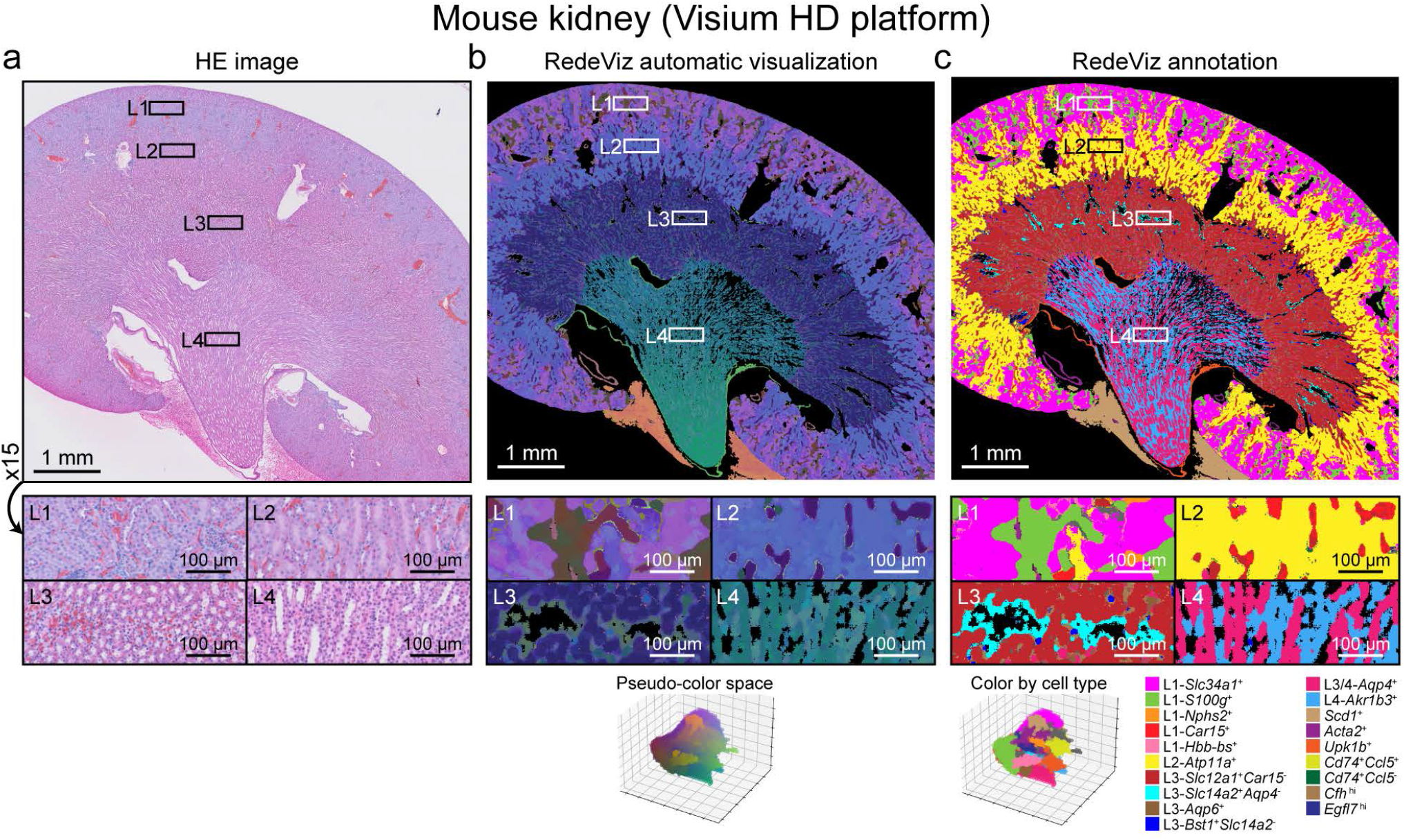
RedeViz visualization on mouse kidney Visium HD dataset. (a) HE image of mouse kidney Visium HD dataset. (b-c) RedeViz automatic cell state visualization (b), and RedeViz annotation (c) after RedeViz enhancement. The images of RedeViz automatic cell state visualization are colored by pseudo-color, and the images of RedeViz cell type annotation are colored by cell types.

### Targeted-enhancement enables computational anatomy iteratively

Finally, we used the Stereo-seq mouse embryo dataset^11^ to evaluate RedeViz’s capability on ST data with organism-level complexity. Using the self-enhancement strategy, RedeViz allows for automatic visualization of embryonic organs, tissues, and cellular contextures with higher resolution compared to the reference annotations without prior knowledge (Figure 6a-b). Various tissues and organs, such as the heart and kidney, are distinctly visible (Figure 6b). Clustering of RedeViz visualization by the Leiden algorithm automatically reveals the structures of heart, liver, lung, kidney, adrenal gland, and GI tract (Figure 6c), consistent with the reference annotations. Regions of the brain, eyes, inner ear, and jaw are even annotated with finer structures. This capacity of RedeViz to enhance ST data and visualize organs/tissues automatically without prior knowledge is especially valuable for highly complex samples or samples from new species, where obtaining a comprehensively annotated reference dataset is particularly challenging.

**Figure 6.**
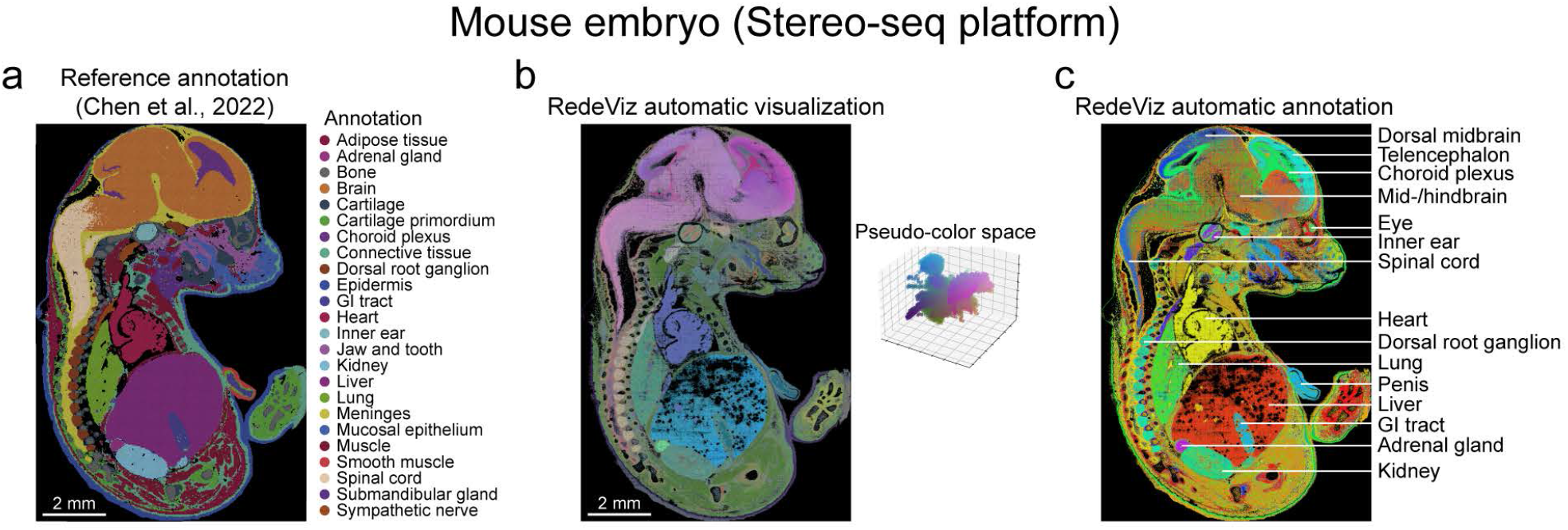
RedeViz visualization on mouse embryo Stereo-seq dataset. (a) Reference annotation of the mouse embryo Stereo-seq dataset. (b) RedeViz automatic cell state visualization after self-enhancement. The image is colored by pseudo-color. (c) RedeViz automatic clustering after self-enhancement. The major organs identified by RedeViz are labeled on the right side.

The visualization of the whole embryo primarily uses colors to discriminate various organs at the organism level during automatic visualization, which is insufficient to visually discriminate different structures within an organ or tissue. To visualize the finer structures within organs or tissues, RedeViz enables a targeted-enhancement strategy. Compared to the self-enhancement strategy, which uses the whole ST dataset as a reference for enhancement, the targeted-enhancement strategy uses the raw signal within the Region of Interest (ROI) instead of the entire dataset as a reference. In the kidney region, the reference annotations categorize this area as kidney and adrenal glands (Figure 7a). While RedeViz’s automatic visualization with the self-enhancement strategy displays detailed structures (Figure 7b), RedeViz with targeted-enhancement revealed finer structural distinctions within the kidneys and the adrenal glands (Figure 7c). Furthermore, fine structures, such as *Podxl*^*+*^ *Cdkn1c*^*+*^ ducts with a 100 μm diameter, were now effectively displayed within the kidneys (Figure 7c) and supported by marker genes (Figure 7d). Similar clear outcomes were achieved for the inner ear regions (Supplementary Figure 7). This strategy proves effective in systematically dissecting the intricate organizational structures of organs and tissues iteratively, offering new avenues for developmental biology and disease research.

**Figure 7.**
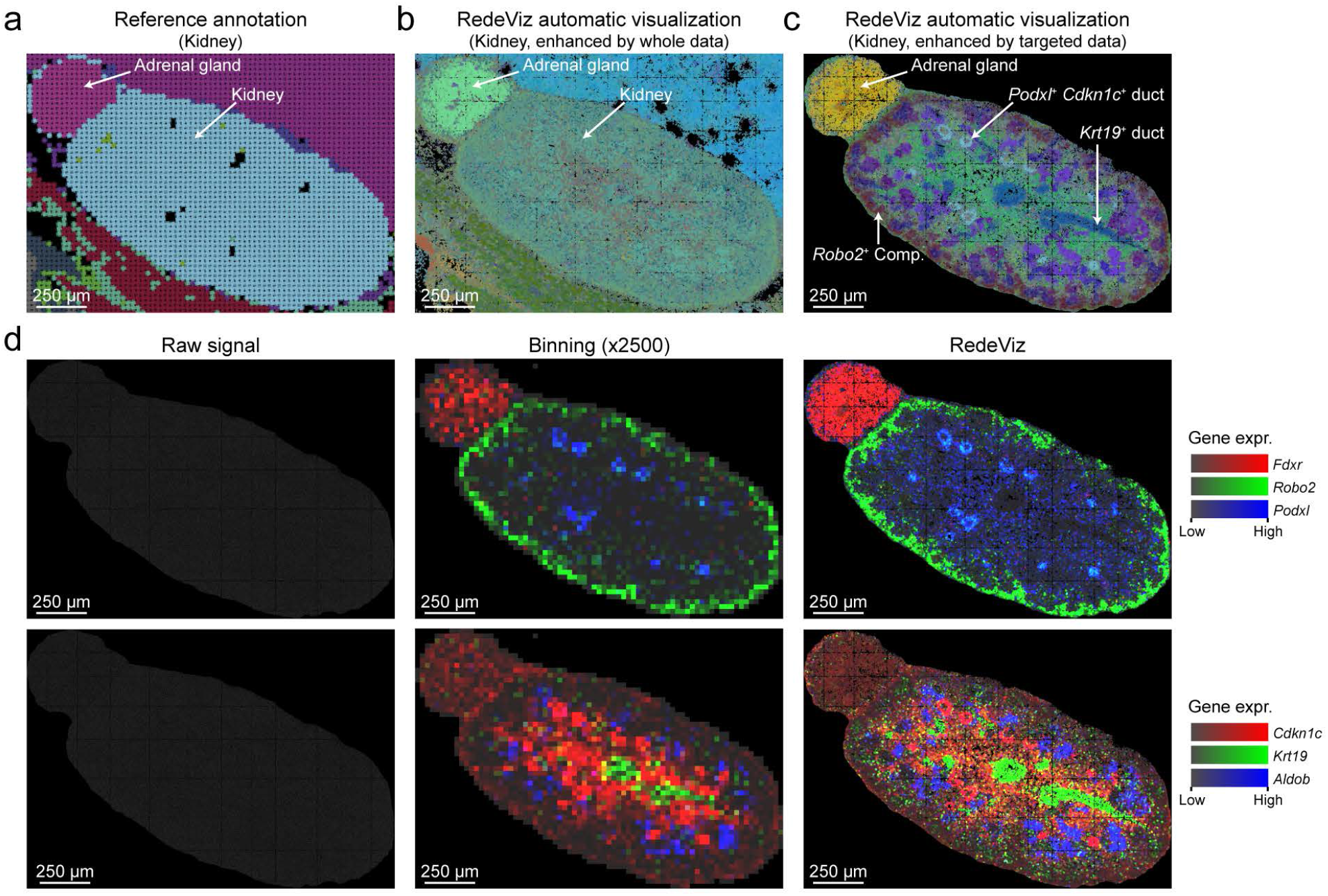
RedeViz targeted-enhancement results of kidney region on the mouse embryo Stereo-seq dataset. (a-c) The reference annotation (a), and RedeViz automatic cell state visualization (b) on the kidney region. (c) RedeViz automatic cell state visualization after targeted self-enhancement on the kidney region. The image is colored by pseudo-color, and structures are labeled on the right side. (d) The Stereo-seq raw signal (left, 500 nm x 500 nm resolution), Stereo-seq binning signal (middle, 25 μm x 25 μm resolution) and RedeViz imputed signal (right, 500 nm x 500 nm resolution) for three marker genes.

## Discussion

In this study, we introduced RedeViz, a segmented-free approach designed to enhance and visualize ST data at the pixel level. RedeViz utilizes diverse strategies to construct references (e.g., scRNA-seq datasets or pre-segmented ST data), to enhance the ST data by dimension raising, to automatically visualize cell states, gene expression patterns, and to illustrate cell contextures. By aligning ST data with reference datasets pixel by pixel, RedeViz resolves data sparsity issues, thereby improving visualizing resolution and the interpretability of spatial patterns. In particular, with whole-transcriptome scRNA-seq data as reference, RedeViz enables imputation of missing genes by ST technologies and clearer spatial patterns by using reference information.

RedeViz demonstrates robust performance across multiple datasets with varying organ/tissue complexity levels. At the tissue-level, it achieves high-resolution visualizations akin to traditional histological images (HE and IF images), especially for cells with complex shapes like adipose tissue and blood vessels. At the organ-level, RedeViz accurately visualizes structures consistent with the cell texture of HE images, as well as visualize smooth muscle-epithelial layer structure with fine details down to a width of 20 μm. For organism-level ST data, RedeViz automatically visualizes entire organs with embryo datasets. Targeted enhancement enables iterative computational anatomy analysis.

The versatility of RedeViz also extends across different types of ST platforms, encompassing both imaging-based methods (such as Xenium, MERFISH, and CosMx) and NGS-based methods (such as Visium HD and Stereo-seq). While RedeViz performs well across most datasets, the performance can be further improved by platform-specific normalization techniques, such as percentile-based gene expression metrics instead of raw expression values to calculate similarity.

Such customized adjustment could help mitigate batch effects between the ST and reference data and thus further improve the performance.

RedeViz still achieves good visualization effects on SP data. However, because genes in SP data are described in terms of relative expression levels, the expression intensities of different genes cannot be directly compared. Therefore, SP data can only be visualized using a self-enhancement strategy for the time being. Additionally, noise signals in SP data, caused by smearing or glare, significantly impact the visualization outcomes. In the future, the development of mapping algorithms between SP and transcriptome data, as well as denoising algorithms for SP data, will further help RedeViz expand its scope of application.

Ultimately, leveraging RedeViz’s WYSIWYG effect for visualization of ST datasets, visual analysis of ST data becomes feasible. In particular, the targeted enhancement strategy may enable computational anatomy analyses based on ST data, which enables deeper insights into the complex structures and processes of biological systems.

## Methods

### RedeViz workflow

The RedeViz algorithm can be subdivided into two fundamental steps: “Raising dimension for enhancement” and “Visualization of enhanced ST data”. The former encompasses three sub-steps: “Deriving information from scRNA-seq data”, “Aligning ST data with scRNA-seq data”, and “Enhancing ST data through dimension raising”. Meanwhile, the latter step manifests in two distinct forms: “Fine-grained visualization” and “Coarse-grained visualization”.

### Deriving information from scRNA-seq data

RedeViz utilizes scRNA-seq data to estimate the derived information including cell states, cell types, and gene expression patterns. Initially, the raw scRNA-seq data is processed by removing genes in the blacklist, including mitochondrial genes, to ensure that subsequent analyses are not influenced by unwanted genes. Cells with individual gene expression accounting for more than 20% of the total gene expressions are filtered out to avoid huge bias from individual genes. Highly variable genes (HVGs) are then identified using the highly_variable_genes function from the scanpy (v 1.9.1) package^15^. And cell coordinates in the phenotypic space are calculated using various embedding methods, such as tSNE-2D, UMAP-2D, and UMAP-3D, based on the expression levels of HVGs. After that, the distribution of gene expression and cell types in the phenotypic space (*PE*_*p*_ and *PC*_*p*_) are estimated based on the average gene expression level and major cell type of neighboring cells. Finally, within the RedeViz framework, each phenotypic coordinate *p* is treated as a cell state, and both the cell state and its corresponding cell type *PC*_*p*_ and gene expression pattern *PE*_*p*_ are regarded as derived information.

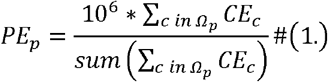

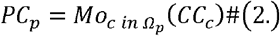

Here, *G* represents the set of genes present in the filtered scRNA-seq data, and *Ω*_*p*_ represents the set of cells that lie within the neighborhood of a given phenotypic coordinate *p. PE*_*p*_ is the estimated phenotypic expression (PE) level of all genes in phenotypic coordinate *p, CE*_*c*_ is the cell expression (CE) levels described as UMI counts of all genes in cell *c. PC*_*p*_ denotes the phenotypic cell type (PC) of phenotypic coordinate *p, CC*_*c*_ denotes the cellular cell type (CC) of cell *c*, 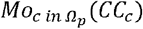 denotes the major cell types of cells in the neighborhood of *p*.

### Aligning ST data with scRNA-seq data

During this step, RedeViz computes the prior probability distribution of similarity between ST data and the cell states derived from scRNA-seq, subsequently mapping the ST data onto the phenotype space established by scRNA-seq.

Initially, RedeViz computes the prior probability distribution. Considering that not all locations in the ST data can be described by the reference phenotypic space, RedeViz introduces two special phenotypic states: ‘background’ for locations with no detectable signal and ‘outer’ for locations that correspond to cell states outside the reference phenotypic space. To estimate the phenotypic expression level of these states, RedeViz calculates the mean value of all PE values and uses this as the background phenotypic expression level of background (*PE*_*bg*_) and outlier (*PE*_*outer*_) expression.

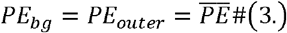

RedeViz makes the assumption that the gene expression levels of ST data 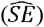 follows a Dirichlet-Multinomial distribution. For each reference phenotype *TruePhe* RedeViz simulates the expression distribution of ST data, and then calculates the cosine similarity score between the simulated ST data and all reference phenotypes plus, including the two special phenotypes. And the distribution of the maximum cosine similarity phenotypes (*ArgMaxPhe*_*TruePhe*_) can be computed by using the Monte Carlo method.

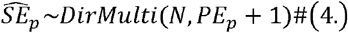

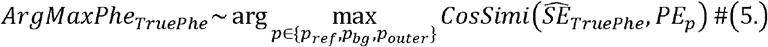

To evaluate whether the ST data is contained within the reference space, RedeViz calculates the distribution of cosine similarity score for each reference phenotype *p* by Monte Carlo method as

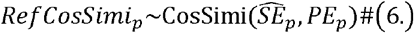

After completing the prior probability distribution calculation, RedeViz projects each pixel in the ST data to the phenotypic space for alignment between ST data and cell states. Similar to the pre-processing of scRNA-seq data, ST genes in the blacklist or whose expression is level higher than 20% of the total UMI are first filtered out. Next, a multiscale average smoothing approach is applied to the spatial expression distribution of each gene. This helps to reduce noise and improve the accuracy of the expression profile at each pixel. Then, RedeViz uses four scores to describe the fitness between ST data and phenotypic projection as following:

Similarity score (*S*_*simi*_):

For a specific location, the similarity score is calculated as the probability that the phenotype *p* is close to the greatest cosine similarity phenotype(*ObsArgMaxPhe*) in statical space. This score measures the gene expression similarity between ST data and phenotypic space.

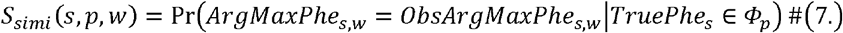

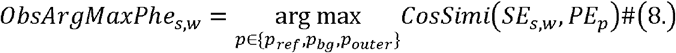

Where *Φ*_*p*_, is the set of phenotypes in the neighborhood of phenotypic coordinate *p,SE*_*SW*_ is the smoothed spatial expression (SE) level at spatial coordinate with window size equal to *w,ObsArgMaxPhe*_*s,w*_ is the phenotype with the greatest cosine similarity to *SE*_*s,w*._

Outer score (*S*_*outer*_):

The outer score is calculated as the normalized distance between the observed and excepted cosine similarity score.

This score measures the tendency of ST data outside the reference phenotypic space.

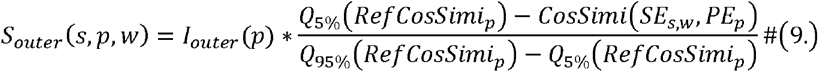

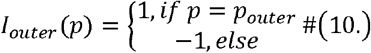

Where *Q*_*x*%_ (*RefCosSimi*_*p*_) is the x% quantile of distribution of *RefCosSimi*_*p*_, and *I*_*outer*_ (*p*) is an indicative function, which is 1 for ‘outer’ phenotype, and -1 for others.

Backround score(*S*_*bg*_):

The background score is calculated as the distance between the specific location signal coverage (*Cov*) and the average signal coverage in the non-zero region (*AveCov*). This score measures the tendency of ST data is in the background area.

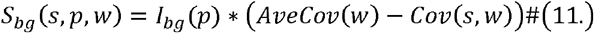

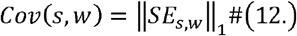

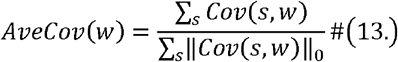

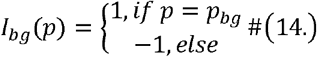

Where *Cov*(*s,w*) is the smoothed total signal coverage at spatial coordinate *s* with window size equal to *w, AveCov*(*w*) is average non-zero signal coverage in ST data, and *I*_*bg*_ (*p*) is an indicative function, which is 1 for ‘background’ phenotype, and -1 for others.

Adjacency score (*S*_*adj*_):

The adjacency score is calculated as the proportion of phenotypes around location in location space close to the phenotype at location in phenotypic space. This score measures how well a location fits in its surrounding phenotypes.

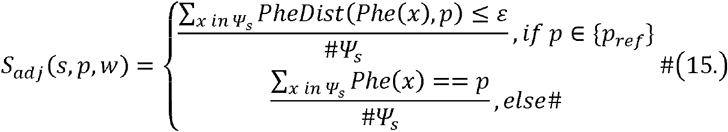

Where *Ψ*_*s*_ is the set of locations adjacent location *s*, and *Phe*(*x*)is the phenotype at location *x*.

RedeViz uses a weighted score (*S*) to measure the projected fitness between ST data and phenotypic space. The phenotypic combination with the highest weighted score is used as the optimized phenotypic projection result (*P*_*opt*_).And the final phenotypic projection (*P*_*final*_) is obtained by performing median smoothing on *P*_*opt*_.

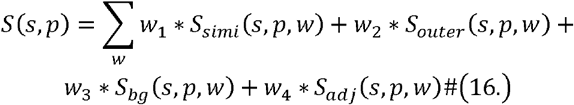

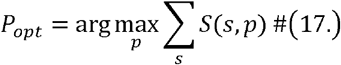

### Enhancing ST data through dimension raising

The original ST data undergoes enhancement through the inclusion of three derived dimensions (dDims): cell state, cell type, and imputed gene expression pattern at measurement resolution. The final phenotypic projection *P*_*final*_ and its corresponding cell type distribution *PC*(*P*_*final*_)are treated as enhancements of the cell state and cell type dimensions, respectively. Regarding the gene expression pattern, RedeViz employs a multi-step process. Initially, Gaussian smoothing is applied to the gene expression within the phenotypic space for a given gene. Subsequently, the gene expression within the phenotypic space is projected onto the ST data, the initial gene expression imputation at measurement resolution is generated. Finally, Gaussian smoothing is carried out on the initial result within the positional space, ultimately yielding the conclusive imputed gene expression pattern.

### Fine-grained visualization

Following the enhancement process, RedeViz becomes capable of visualizing the distributions of cell states, cell types, and gene expression patterns at measurement resolution. Through the translation of derived cell state coordinates into RGB coordinates for every pixel, RedeViz implements automatic cell state visualization. Additionally, by assigning custom colors to either cell states or cell types, RedeViz enables customized visualization of cell states or cell types. And by converting gene expression levels into intensities, RedeViz performs the visualization of genes expression patterns unmeasured by ST data.

### Coarse-grained visualization

RedeViz can identify and visualize multi-scale spatial domains based on the enhanced ST data. In this project, we conceptualize spatial domains as continuous regions with similar cell states. If we shift our perspective from the phenotype space to the color space, the problem of spatial domain segmentation aligns naturally with image segmentation. Therefore, RedeViz adopts a two-step approach to address this problem. RedeViz first performs Gaussian blur for each phenotypic dimension of the derived cell state projection. Then, the meanShiftSegmentation function in OpenCV (v4.5.2) package is used for plane segmentation. By adjusting the thresholds of spatial distance (which relates to the smooth window), color distance (which relates to the cell type purity), and segment size (which relates to the domain size) within the meanShiftSegmentation function, RedeViz can divide the ST data into domains of different granularities from mesoscope to macroscope.

### Automatic self-enhanced visualization and targeted-enhanced visualization

RedeViz can perform self-enhanced visualization using solely ST data. The process begins by extracting the total expression level of the gene. Subsequently, after applying two-dimensional Gaussian filtering, the region where total expression is less than 20% of the median value of non-zero signals is designated as the background area, with the corresponding signal value set to 0. Cell segmentation is achieved through the maskSLIC algorithm, based on the total expression distribution, And the mask region of the maskSLIC algorithm is computed using the adaptiveThreshold function in the OpenCV package. By counting the gene expression signals within each cell segmentation region, a cell-gene expression matrix is generated. Treating this expression matrix as single-cell data, and employing the Leiden algorithm for cell clustering, a reference dataset is created. Finally, by utilizing this pre-segmented reference to enhance ST data, RedeViz is capable of automatic self-enhanced visualization. By selecting a Region-Of-Interest (ROI) from the entire ST data and performing self-enhanced visualization within the ROI, we can achieve targeted-enhanced visualization.

### Processing of breast cancer Xenium dataset

The reference scRNA-seq data was annotated using scanpy pipeline. In detail, cells with gene number less than 500 or mitochondrial gene ratio greater than 15 were first filtered out from the reference single-cell data. After normalization, the highly variable genes (HVGs) were identified by highly_variable_genes function. Based on the HVGs, a nearest neighbor graph was built and used to visualize the cells in two dimensions through the UMAP algorithm. The Leiden algorithm was then used for unsupervised clustering. To annotate the cell types within each cluster, the rank_genes_groups function was used to identify cluster specific marker genes. And these cluster-specific marker genes were then used to assign cell types to each cluster.

For processing Xenium data, the BLANK genes in the Xenium dataset were removed because they represent background noise and do not contribute to the gene expression patterns of cells. Cells with Xenium gene number less than 20 or Xenium gene UMI less than 100 were filtered out to ensure that only cells with sufficient gene expression information were used in the further analysis. Then the Xenium gene expression levels of pre-segmentation Xenium dataset, filtered 3’-seq and 5’-seq were used as reference data to perform the standard RedeViz workflow to enhance and visualize Xenium dataset.

### Processing of mouse ileum MERFISH and NSCLC CosMx dataset

The standard RedeViz workflow and the RedeViz automatic self-enhanced visualization workflow were used to enhance and visualize MERFISH and CosMx dataset.

### Processing of human bone marrow CODEX dataset

We employed the following method to normalize CODEX signals as gene expression signals for input into RedeViz:

1. Division of DAPI Image: We used a Gaussian process classifier to divide the DAPI image into signal areas and background areas.
2. Calculation of Signal Intensities: For each signal image, we calculated the non-zero median signal intensity (*Q*_*low*_) of the background area and the 99.95% quantile signal intensity (*Q*_*hi*_) of the signal area.
3. Signal Standardization: We standardized the signal using the formula: 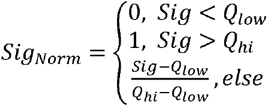

These pre-segmentation outcomes were used as reference data to enhance and visualize the CODEX dataset with RedeViz.

### Processing of mouse kidney Visium HD dataset

The nuclei segmentation of raw Visium HD data was performed with official workflow^10^. And the standard RedeViz automatic self-enhanced visualization workflow was used to enhance and visualize this dataset.

### Processing of mouse embryo Stereo-seq dataset

The pre-segmentation Stereo-seq data was used as reference, and the standard RedeViz workflow was used to enhance and visualize the Stereo-seq dataset. The distribution of cell states were visualized by RedeViz with default parameter, and automatic annotation was clustered by the Leiden algorithm and named according to the anatomical location. Specifically, regions of the kidney and inner ear were selected for targeted-enhanced visualization.

### Data availability

The breast cancer Xenium dataset and mouse kidney Visium HD dataset were downloaded from the 10x Genomics official database^9^. The mouse ileum MERFISH dataset was downloaded from the original publication^12^, and the reference scRNA-seq data was downloaded from single cell portal database (accession ID: SCP1038). The mouse NSCLS CosMx dataset^8^ was downloaded from the NanoString official database. The human bone marrow CODEX dataset was downloaded from FigShare database (DOI: 10.25452/figshare.plus.c.7174914). The mouse embryo Stereo-seq dataset was downloaded from the original publication^11^, including the information of Stereo-seq raw signal, pre-segmented cell-gene expression matrix, and corresponding cell type annotation.

### Code availability

The code of RedeViz is implemented in Python and stored at https://github.com/sqreb/RedeViz. The benchmarking and analysis codes are provided at https://github.com/sqreb/RedeVizMs.

## Acknowledgments

This work was supported by Changping Laboratory, the National Natural Science Foundation of China (32022016 X.R., 92159305 X.R., and 31991171X.R.), National Key R&D Program of China (2020YFE0202200 X.R. and 2022YFC3400904 X.R.).

## Author Contributions

X.R. conceived the study. D.W performed the analysis and wrote the code. D.W and X.R. wrote the manuscript together.

## Supplementary figures and legends

**Supplementary Figure 1. Schematic workflow of the Raising Dimension for Enhanced Visualization (RedeViz) algorithm**. The original sparse ST data undergoes enhancement via alignment with reference data, allowing to incorporate additional derived dimensions (dDim) extracted from scRNA-seq data or pre-segmented ST data, including cell types, cell states, and gene expression signal. Following this enhancement, the fine-grained visualization of cell states, cell types, and imputed gene expression signals at measurement resolution can be directly performed. By adjusting the resolution parameters, RedeViz can also identify and visualize multi-scale spatial domains with biological functions at a coarse-grained level using the enhanced ST data. The comprehensive visualization of information spanning diverse dimensions and levels of detail within the ST data, allowing RedeViz to closer to realize the “What You See Is What You Get” (WYSIWYG) effect.

**Supplementary Figure 2. Gene expression imputation on breast cancer Xenium dataset**. (a-b) HE image (a) and the total Xenium signal distribution. (b-d) The gene expression pattern of Xenium raw signal (b, 500 nm x 500 nm resolution), Xenium binning signal (c, 25 μm x 25 μm resolution) and RedeViz imputed signal (d, 500 nm x 500 nm resolution) of two Xenium included genes. (e) RedeViz gene expression imputation of two Xenium excluded genes.

**Supplementary Figure 3. RedeViz visualization on mouse ileum MERFISH dataset enhanced by scRNA-seq**. (a-b) RedeViz automatic cell states visualization (a) and cell type clustering (b) enhanced by scRNA-seq. The images of RedeViz automatic cell state visualization colored by pseudo-color. (e) RedeViz gene expression imputation for three marker genes (*Alpi*: Enterocyte cells; *Mki67*: TA cells; *Gm15293*: Paneth cells).

**Supplementary Figure 4. Automatic visualization and gene expression imputation on human non-small cell lung cancer (NSCLC) CosMx dataset**. (a) RedeViz cell type visualization. The image is colored by pseudo-color. (b-d) The RedeViz imputed signal (b, 500 nm x 500 nm resolution), CosMx signal (c, 500 nm x 500 nm resolution), CosMx binning signal (d, 25 μm x 25 μm resolution) of three marker genes (*CD163*: Macrophage cells; *AGR2*: Tumor cells; *MS4A1*: B cells).

**Supplementary Figure 5. CODEX signal of human bone marrow CODEX dataset**. CODEX signal of cell type specific proteins, related to figure 4d.

**Supplementary Figure 6. Gene expression imputation on mouse kidney Visium HD dataset**. The specific cell state visualization (left, 500 nm x 500 nm resolution) and distribution of corresponding marker genes (right, 10 μm x 10 μm resolution) of 12 annotated cell types.

**Supplementary Figure 7. RedeViz targeted-enhancement results of inner ear region on mouse embryo Stereo-seq dataset**. (a-c) The reference annotation (a), and RedeViz automatic cell state visualization (b) on the inner ear region. (c) RedeViz automatic cell state visualization after targeted self-enhancement on the inner ear region. The image is colored by pseudo-color, and structures are labeled on the right side. (d) The Stereo-seq raw signal (left, 500 nm x 500 nm resolution), Stereo-seq binning signal (middle, 25 μm x 25 μm resolution) and RedeViz imputed signal (right, 500 nm x 500 nm resolution) for three marker genes.

